# CD20 as a gatekeeper of the resting stage of human B cells

**DOI:** 10.1101/2020.07.31.230110

**Authors:** Kathrin Kläsener, Julia Jellusova, Geoffroy Andrieux, Ulrich Salzer, Chiara Böhler, Sebastian N. Steiner, Jonas B. Albinus, Marco Cavallari, Beatrix Süß, Melanie Boerries, Reinhard E. Voll, Bernd Wollscheid, Michael Reth

**Affiliations:** Biology III, faculty of biology, University Freiburg; Research Centres BIOSS and CIBSS; Institute of Medical Bioinformatics and Systems Medicine, Medical Center – University of Freiburg, Faculty of Medicine, University of Freiburg, Freiburg 79110, Germany; German Cancer Consortium (DKTK) Partner Site Freiburg, German Cancer Research Center (DKFZ), Heidelberg, Germany; Department of Rheumatology and Clinical Immunology, Medical Center – University of Freiburg, Faculty of Medicine, University of Freiburg, Freiburg, Germany; Institute of Translational Medicine at the Department of Health Sciences and Technology, ETH Zurich, 8093 Zurich, Switzerland; Swiss Institute of Bioinformatics (SIB), Lausanne, Switzerland; Department of Biology, Centre for Synthetic Biology, Technical University Darmstadt, Darmstadt, Germany; Comprehensive Cancer Center Freiburg (CCCF), Medical Center – University of Freiburg, University of Freiburg 79106, Germany

## Abstract

CD20 is a B cell specific membrane protein and a target of therapeutic antibodies such as rituximab (RTX)^1^. In spite of the prominent usage of anti-CD20 antibodies in the clinic little is known about the biological function of CD20^2^. Here we show that CD20 controls the nanoscale organization of receptors on the surface of resting B lymphocytes. A CRISPR/Cas-based ablation of CD20 in Ramos B cells results in a relocalisation of the IgM B cell antigen receptor (IgM-BCR) and the co-receptor CD19. The resulting IgM-BCR/CD19 signaling synapse leads to transient B cell activation followed by plasma cell differentiation. Similarly to CD20-deficient Ramos cells, naïve human B cells treated with rituximab in vitro or isolated from patients during rituximab administration display hallmarks of transient activation characterized by the formation of the IgM-BCR/CD19 signaling synapse, followed by CD19 and IgM-BCR downregulation. Moreover, increased expression of specific plasma cell genes can be observed after rituximab treatment in relapsed CLL patients. In summary we identify CD20 as a gatekeeper of the resting state on human B cells and demonstrate that a disruption of the nanoscale organization of the B cell surface via CD20 deletion or anti-CD20 treatment profoundly alters B cell fate.

---

The human Burkitt lymphoma cell line Ramos carries large amounts of the IgM-BCR on its cell surface, whereas the IgD-BCR is less abundant on these cells (**Fig 1a**). Ramos B cells also express the co-receptor CD19 and the B cell-specific membrane protein CD20^3–5^. The B cellspecific expression of CD20 makes it an ideal target for therapeutic antibodies used for the treatment of human B cell neoplasias and autoimmune diseases such as rheumatoid arthritis or systemic lupus erythematosus^6–8^. We have previously localized the CD19/CD81 coreceptor module together with CD20 and CD40 inside an IgD-class specific membrane compartment on the surface of resting B cells^9–11^. To determine the nanoscale organization of the CD20 molecule on the Ramos B cell surface, we used the Fab-based proximity ligation assay (Fab-PLA). Similar to our previous studies on primary B cells, we find CD20 in close proximity to the IgD-BCR on resting Ramos cells whereas after a pervanadate-mediated B cell activation CD20 moves to the IgM-BCR (**Fig 1b**). Thus, the nanoscale membrane organization of CD20 seems to be similar on Ramos and primary B cells (**Fig 1c**).

**1.**
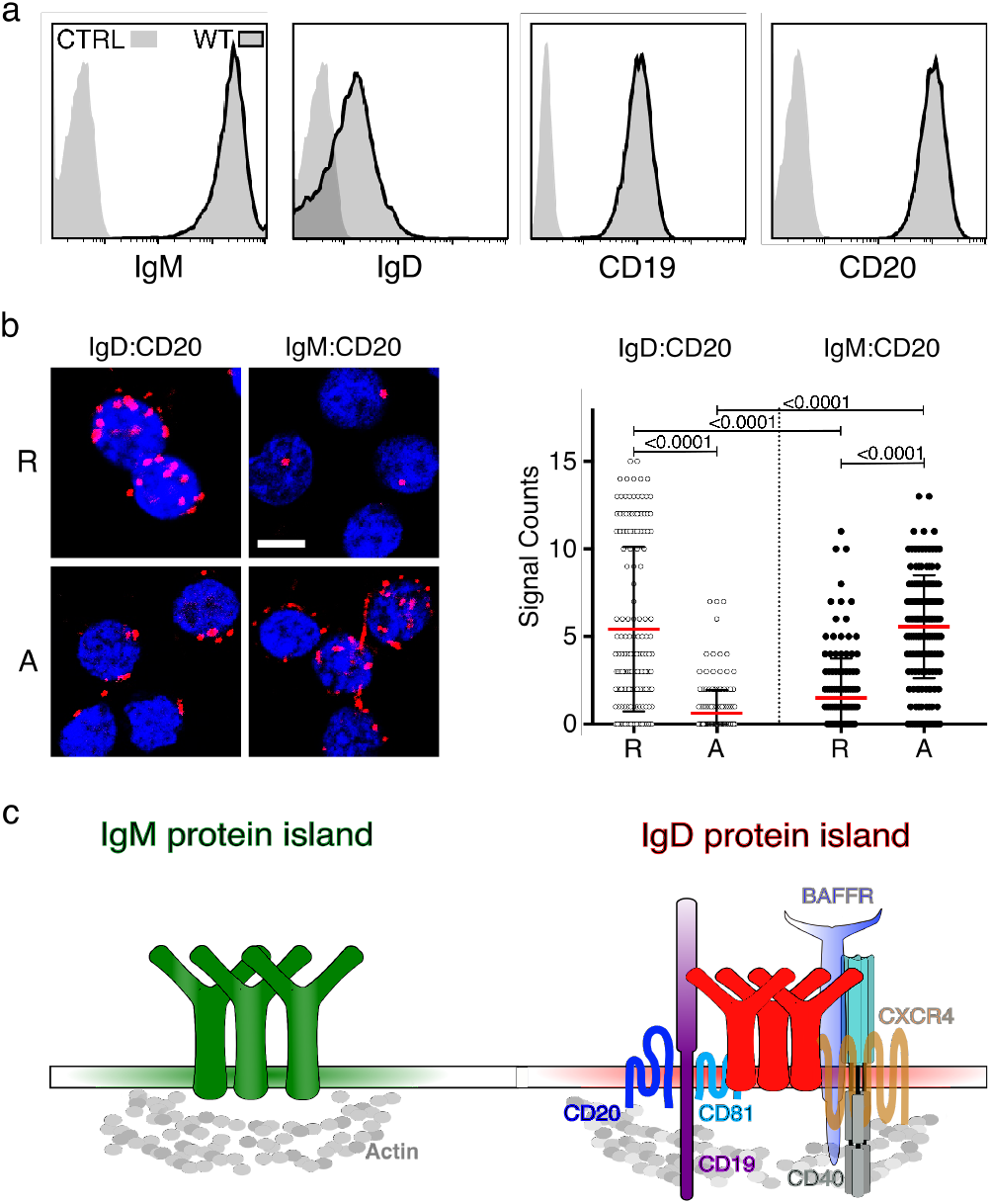
CD20 expression and localization on WT Ramos B cells. **a** Flow cytometry analysis showing the expression of IgM-BCR, IgD-BCR, CD19 and CD20 on Ramos WT cells compared to unstained control (CTRL), n=18 **b** Fab-PLA of the proximity of CD20 to IgD-BCR or CD20 to IgM-BCR on resting (R), or 5 min pervanadate activated (A) WT Ramos B cells. Representative microscope images (left) of PLA-signals shown in red and nuclei in blue. Scale bar 10 μM. Scatter dot plot represent the mean (red bar) and SD of PLA signals (Signal Counts), n=3 **c** Schematic drawing of the proposed model of IgM and IgD-BCR protein islands with coreceptors on the surface of human resting B cells.

Using the CRISPR/Cas9 technology we targeted exon 3 of the MS4A1 gene encoding CD20 and disrupted the open reading frame of this gene shortly after the ATG start codon (**Extended Data Figure 1**). The obtained CD20KO Ramos cell lines were classified according to the date of gene targeting as new (KO-N), intermediate (KO-I), and late (KO-L) cells (**Fig 2a**). The KOI and KO-L cells were negatively selected for the loss of CD20 expression by cell sorting. In total we generated 18 distinct CD20KO Ramos cell lines (**Supplementary Table 5**) that were analysed at different time points for the abundance of selected cell surface markers. Surprisingly, the KO-I and KO-L Ramos cells not only lost the surface expression of CD20, but also that of other B cell surface markers such as CD19, CD22, CD81, CD40 and the IgM-BCR (**Fig 2b**). The altered protein abundance was confirmed by Western blotting analysis of lysates from KO-L cells which showed a loss, or marked reduction of CD20, CD19, CD22, CD40, IgM and CD81 proteins in comparison to Ramos wild type (WT) cells (**Fig 2c**). Thus, the CRISPR/Cas9-induced CD20 gene deficiency is accompanied by drastic changes of the receptor abundances on the surface of Ramos cells and the loss of the B cell resting stage.

**2.**
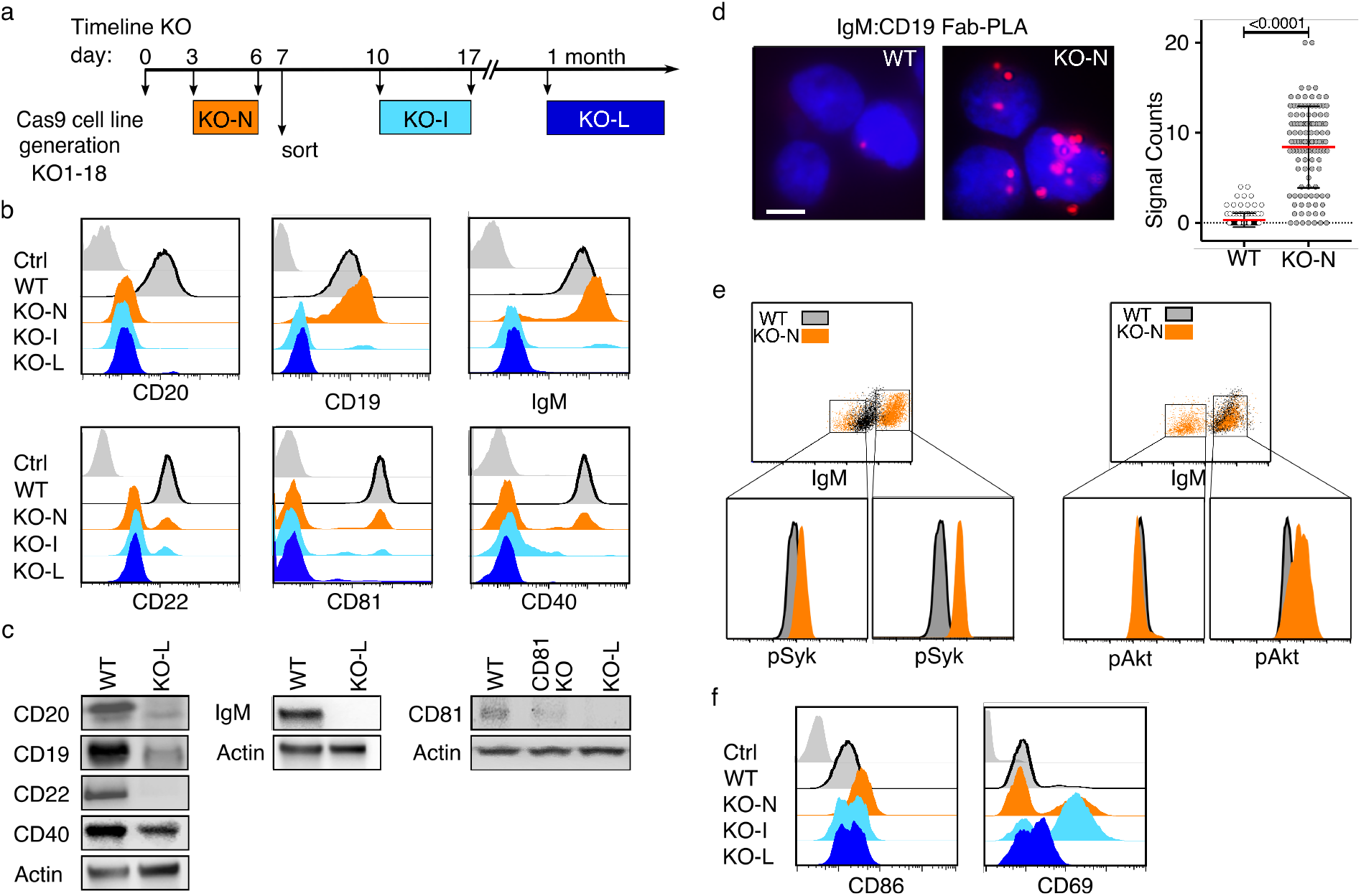
Loss of CD20 results in altered expression of B cell surface markers and transient activation. **a** Timeline of developmental states in CD20KO Ramos cell-line generation. MS1A4 gene targeting with CRISPR/Cas9 at day 0, developmental stage at day 3-6 shown as CD20KO-new (KO-N, orange), at day 10-17 shown as CD20KO-intermediates (KO-I, light blue), after 1 month shown as CD20KO-late (KO-L, dark blue). **b** Flow cytometry analysis showing overtime expression of surface molecules of KO-N, KO-I, and KO-L Ramos cells compared to WT and unstained control (grey), n=18. **c** Representative western blot analysis of KO-L Ramos B cell-lysates (right) compared to WT (left). Lysates were taken 20 days after MS4A1 gene targeting, n=14 **d** Fab-PLA analysis of IgM-BCR proximity to CD19 in resting WT Ramos B cells compared to the unstimulated KO-N cells 3 days after MS4A1 gene targeting or with empty vector (not shown), Scale bar 10μM (left). PLA microscope images were quantified as scatter dot plot with mean and SD (right), n=3 **e** Intracellular flow cytometry analysis of phosphorylated Akt (pAkt-Ser473) or phosphorylated Syk (pSyk-Tyr525,526) of KO-N cells compared to WT. Gating used to analyse high IgM-BCR expressing KO-N cell population for pSyk or pAkt levels respectively. **f** Flow cytometry analysis showing the overtime expression of the B cell-specific surface activation markers CD86, and CD69 of KO-N compared to WT.

## Loss of CD20 expression is associated with B cell activation

Similarly to KO-I and KO-L cells, KO-N Ramos cells show reduced expression of CD20, CD22, CD81 and CD40, however CD19 and IgM-BCR levels are increased (**Fig 2b**). On resting murine and human B cells, CD19 and the IgM-BCR are separated from each other but form a signaling synapse upon B cell activation^9,12^. Interestingly, an increased IgM:CD19 proximity is detected by Fab-PLA on KO-N Ramos cells, indicating that these cells are activated (**Fig 2d**). Indeed, a phosphoflow cytometry analysis of WT and KO-N Ramos cells revealed an increased phosphorylation of Syk and Akt in the KO-N population with the highest IgM expression (**Fig 2e**). Furthermore, KO-N cells upregulate the B cell activation markers CD86 and CD69 (**Fig 2f**). Thus, the CD20 gene deletion is associated with a transient B cell activation followed by a loss/downregulation of several B cell surface markers. Together, these data suggest that on resting B cells CD20 functions as a gatekeeper preventing uncontrolled IgM-BCR/CD19 interaction and signaling.

To learn more about the molecular requirements for the changes of B cell surface expression displayed by the CD20KO Ramos cells we generated CD20 loss mutants from Ramos CD19KO and from BCR-KO cells, deficient in heavy chain, light chain and activation induced deaminase (AID) expression. Interestingly, none of the double KO (DKO) Ramos cells displayed the transient B cell activation (data not shown). Unlike KO-L, the CD19/20DKO-L Ramos cells still express the IgM-BCR and CD22 (**Extended Data Figure 2a**), whereas the BCR/20DKO-L Ramos cells retain expression of the B cell surface marker CD19 and CD22 (**Extended Data Figure 2b**). Together this analysis suggests that the BCR and CD19 interaction is indispensable for the transient B cell activation and the altered surface expression of CD20KO Ramos B cells.

## Re-expression of CD20 restores the resting B cell phenotype

We next asked whether the re-expression of CD20 could restore the expression of cell surface proteins on CD20-deficient Ramos B cells. For this we established a conditional KO (cKO) of the CD20 encoding MS4A1 gene (**Extended Data Figure 3**). Using the CRISPR/Cas9 technique, we inserted between exon 3 and exon 4 of the MS4A1 gene, an aptamer-controlled conditional exon (c-exon) whose incorporation in the processed MS4A1 mRNA results in a stop code thus rendering this transcript defective^13^. Upon exposure of the Ramos cKO cells to tetracycline (Tet) the antibiotic molecule binds to and stabilizes an aptamer whose sequence partly overlaps with the 3’ splice site of the c-exon thus preventing its incorporation into the CD20 transcript (**Extended Data Figure 3a**). This synthetic tetracycline riboswitch enables the successful regulation of CD20 protein levels. Similar to KO-L, the cKO Ramos B cells cultured for more than 30 days do not express CD20 nor CD19, IgM-BCR, CD22, CD81 or CD40 on the B cell surface (**Extended Data Figure 3b**). An overnight (o/n) Tet-exposure of the cKO Ramos cells restores the expression of CD20 and that of the other B cell markers on the cell surface. Thus, upon its re-expression, CD20 can resume its gatekeeper function to support the resting state of Ramos B cells.

## Rituximab treatment of naïve human B cells terminates the gatekeeper function of CD20

CD20 is a prominent target of therapeutic antibodies such as rituximab (RTX) that are used for the depletion of B lymphocytes in autoimmune diseases and B cell neoplasias. To test whether the RTX treatment interferes with the gatekeeper function of CD20 we used flow cytometry to monitor the expression of B cell surface markers on WT Ramos cells (**Fig 3a**), and healthy donor (HD) derived naïve B cells (**Fig 3b)** before or after a RTX treatment. Interestingly, all RTX-treated cells lost CD20 from the surface and simultaneously downregulated CD19, CD22 and the IgM-BCR to a similar extent as the KO-L Ramos cells. To test for the immediate effect of RTX *in vivo* we analysed blood samples taken from patients suffering from rheumatoid arthritis (RA) undergoing RTX treatment. B cells from these patients showed a transient increase of surface IgM-BCR expression already after 15 min, followed by a continuous IgM-BCR downregulation. CD19 surface expression, which is routinely used as a B cell marker was almost lost after 30 min of RTX treatment, although B cells were still present and detectable (**Fig 3c**). A Fab-PLA study showed an increased BCR-IgM:CD19, Igα:pSyk, Igα:pYp110δ and Igα:pAkt proximity in RTX-treated compared to untreated HD B cells, indicating that the RTX treatment is accompanied by increases of BCR-associated Syk/PI-3kinase signaling and B cell activation (**Extended Data Figure 4a and 4b**). In summary, these data show that CD20 is a gatekeeper also for resting naïve human B cells and that a CD20 deficiency or RTX treatment terminates B cell dormancy.

**3.**
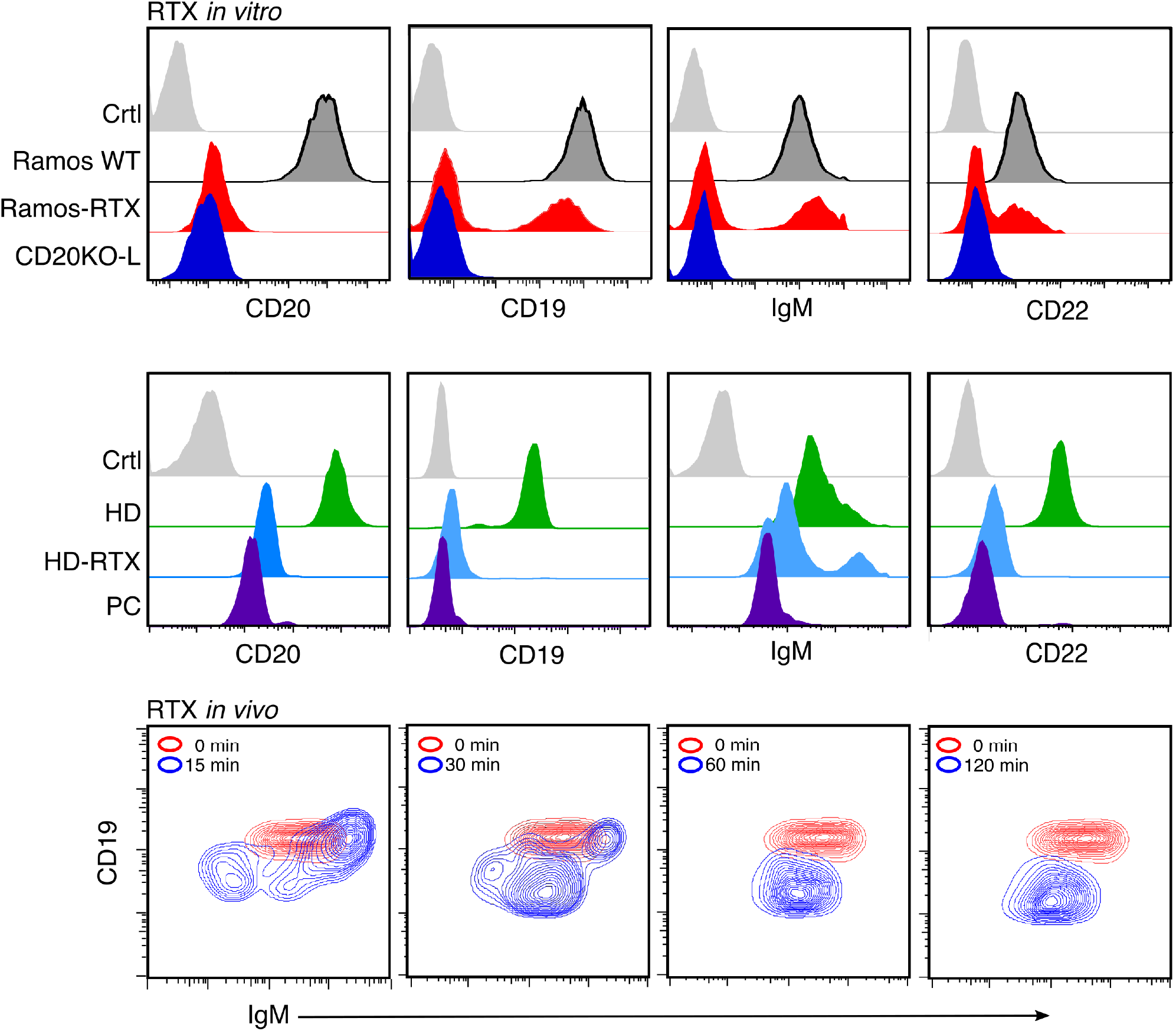
*In vitro* and *in vivo* treatment with rituximab (RTX) leads to CD20KO phenotype. **a** Flow cytometry analysis after treatment with RTX for 3 days showing the loss of CD19, IgM-BCR and CD22 on Ramos-RTX compared to CD20KO-L, untreated Ramos-WT, and unstained control (Ctrl), n=3 **b** Flow cytometry analysis of negatively selected naïve B cells from peripheral blood of a healthy donor were treated with RTX for 60min (HD-RTX) and compared to untreated (HD), to plasma cells (PC) of the same donor or left untreated and unstained (Ctrl) n=3. **c** Example of B cells taken from EDTA whole blood samples of RA patient undergoing RTX treatment after 0, 15, 30, 60, 120 min. RTX [1mg/mL] flow rate 50mL/h. Flow cytometry staining of CD19+/ CD27-/ IgM+ selected B cells shows internalization of IgM and CD19, n=2.

## Dynamic alteration of the surfaceome of CD20-deficient B cells

To obtain a more global picture of the alteration of protein expression on the surface of CD20-deficient Ramos B cells we took advantage of the Cell Surface Capture (CSC) technology^14,15^. CSC technology allows for the efficient labelling, identification and relative quantification of N-glycosylated proteins residing on the cell surface and provides a snapshot of the surfaceome without having to use antibodies to broadly phenotype the B cells^16^. The CSC analysis of the 3-7-day-old KO-N Ramos cells shows a significant upregulation of surface markers (3 down versus 45 up, p-value <0.05) that is consistent with the activated state of these cells (**Extended Data Figure 5a and 5b**). The proteins with higher cell surface abundance include the IgM-BCR and CD19, thereby confirming our previous flow cytometry analysis (**Fig. 2b**). Interestingly, the loss of CD20 on KO-N Ramos cells resulted in the upregulation of several tetraspanins such as CD37, CD53, CD63, CD82, and TSN3 that could contribute to nanoscale membrane organization as well^17^. In contrast to KO-N, the KO-L Ramos cells display a reduction in the cell surface abundance of many proteins (123 down versus 58 up, p-value <0.05), indicating that at a later time point the CD20 deficiency is associated with gene silencing (**Extended Data Figure 5c and 5d**). Indeed, the abundance of many B cell markers such as the BCR, CD19, CD38, CD27 and CD22 is reduced on KO-L Ramos cells. Interestingly, several surface proteins upregulated on KO-L Ramos cells are also found to be highly abundant on the surface of plasma cells. This suggests that the loss of CD20 is not only associated with a loss of the resting B cell stage but also an increased differentiation towards the plasma cell stage.

## Plasma cell differentiation of CD20-deficient B cells

In culture, with time CD20KO Ramos cells become larger as indicated by the increased forward scatter (FSC) and they upregulate TACI and CD138, both plasma cell markers. (**Fig 4a**). An intracellular FACS analysis showed that CD20KO Ramos cells upregulate AID at the early KO-N stage where these cells display active Syk/PI-3 kinase signaling. Thus, similar to activated B cells participating in a germinal center (GC) reaction the CD20KO Ramos seem to undergo somatic hypermutation and class switching. Indeed, the KO-L Ramos cells carry IgA on their cell surface and secrete IgA (**Fig 4b**). In normal activated B cells, the differentiation to the plasma cell stage is associated with a transcriptional switch involving the down-regulation of the transcription factor PAX5 and upregulation of Blimp-1. Western blot analysis shows that this is also the case in CD20KO Ramos cells (**Fig 4c**). In comparison to WT, the KO-L Ramos cells express less PAX5 and more Blimp-1. Furthermore, KO-L Ramos cells contain more FoxO1, IRF4 and XBP1s, transcription factors that are associated with B to plasma cell differentiation. Another known regulator of plasma cell differentiation is the Blimp-1 suppressor and transcriptional silencer Bcl6, that is highly expressed in GC B cells as well as GC-derived B lymphomas such as Ramos^18–21^. Indeed, Bcl6 is expressed in WT Ramos but no longer detectable in the KO-L cells. The PAX5 to Blimp-1 transcriptional switch was not detected in the CD19/20DKO-L and BCR/20DKO-L Ramos cells, indicating that BCR/CD19 signaling and transient B cell activation was required for this *in-vitro* human plasma cell differentiation (**Extended Data Figure 6c and 6d**). We next asked whether a RTX treatment is also accompanied by an increased plasma cell differentiation of B cells *in vivo*. For this we analysed the transcriptome profile of B cells from CLL patients^22^ that have relapsed after a combined RTX treatment (RTX± Fludarabine) for the expression of plasma cell specific genes in a gene set enrichment analysis (GSEA) and compared this gene set with Ramos WT and CD20KO-L cells (**Fig 4e)**. Interestingly, the samples from 11 of 13 CLL-patients showed a significantly increased expression of plasma cell-specific genes after combined RTX-treatment, confirming our results that CD20-derived B cell activation can induce activation of the plasma cell transcription program (**Fig 4f and 4g**).

**4.**
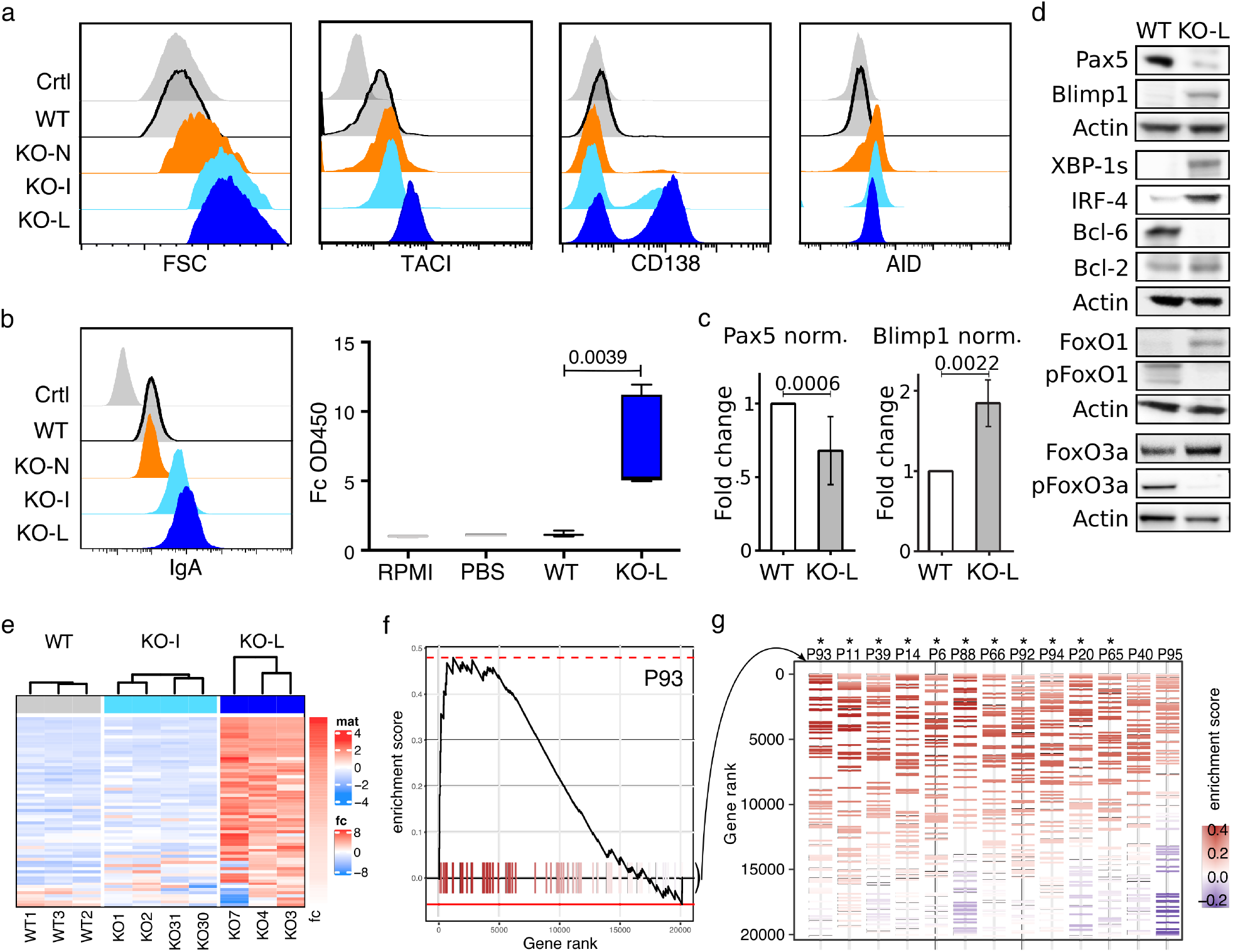
Increased Plasma cell differentiation in CD20KO Ramos B cells. **a** Flow cytometry analysis showing size (FSC-A) and expression of plasma cell markers TACI, CD138, and AID on KO-N, KO-I, and KO-L Ramos B cells compared to WT and unstained control (Ctrl), n= 6. **b** Expression of surface IgA-BCR (left) and ELISA of IgA secretion (right) of KO-L compared to Ramos WT, PBS and RPMI control. **c** Summarized intracellular flow cytometry analysis of 6 independently generated CD20KO Ramos B cell-lines showing the fold change (FC) of Pax5 and Blimp1 expression of KO-L Ramos B cell lines compared to WT. **d** Representative examples of western blot analysis for B cell differentiation markers of KO-L Ramos B cells compared to WT. Lysates were taken 20 days after induction of CD20KO. n=3. **e** PLASMA UP GENES Heatmap, expression of plasma cell differentiation up-regulated genes. Color code indicates the row-wise scaled intensity across the samples. Genes are ranked according to their Log2 fold FC in WT (left) vs. KO-L (right) n≥3. **f** Example of enrichment plot (Curve) of plasma cell differentiation up-regulated genes in one patient. **g** Enrichment barcode illustrating the distribution of plasma cell differentiation up-regulated genes (colored segments) in every individual patient. from GSE37168. Treated patients were ordered from left to right based on their enrichment score, from high to low. Significant enrichment scores are depicted by “*”.

## Distinct alterations of cell metabolism during Ramos plasma cell development

Plasma cells are large cells with an extended endoplasmic reticulum (ER) and a high rate of antibody production. Plasma cells thus have a higher demand for energy and biosynthetic precursor molecules than B lymphocytes^23,24^. To characterize the metabolic program of WT and KO-L Ramos cells we performed metabolic flux experiments and untargeted metabolomics analyses. We found the abundance of 117 biochemicals to be significantly increased and the abundance of 214 biochemicals to be significantly decreased in KO-L Ramos cells in comparison to WT cells. Analysis of the oxygen consumption rate (OCR) showed that KO-L cells consume more oxygen in the basal state and possess a higher spare respiratory capacity than WT Ramos cells (**Fig 5a**).

**5.**
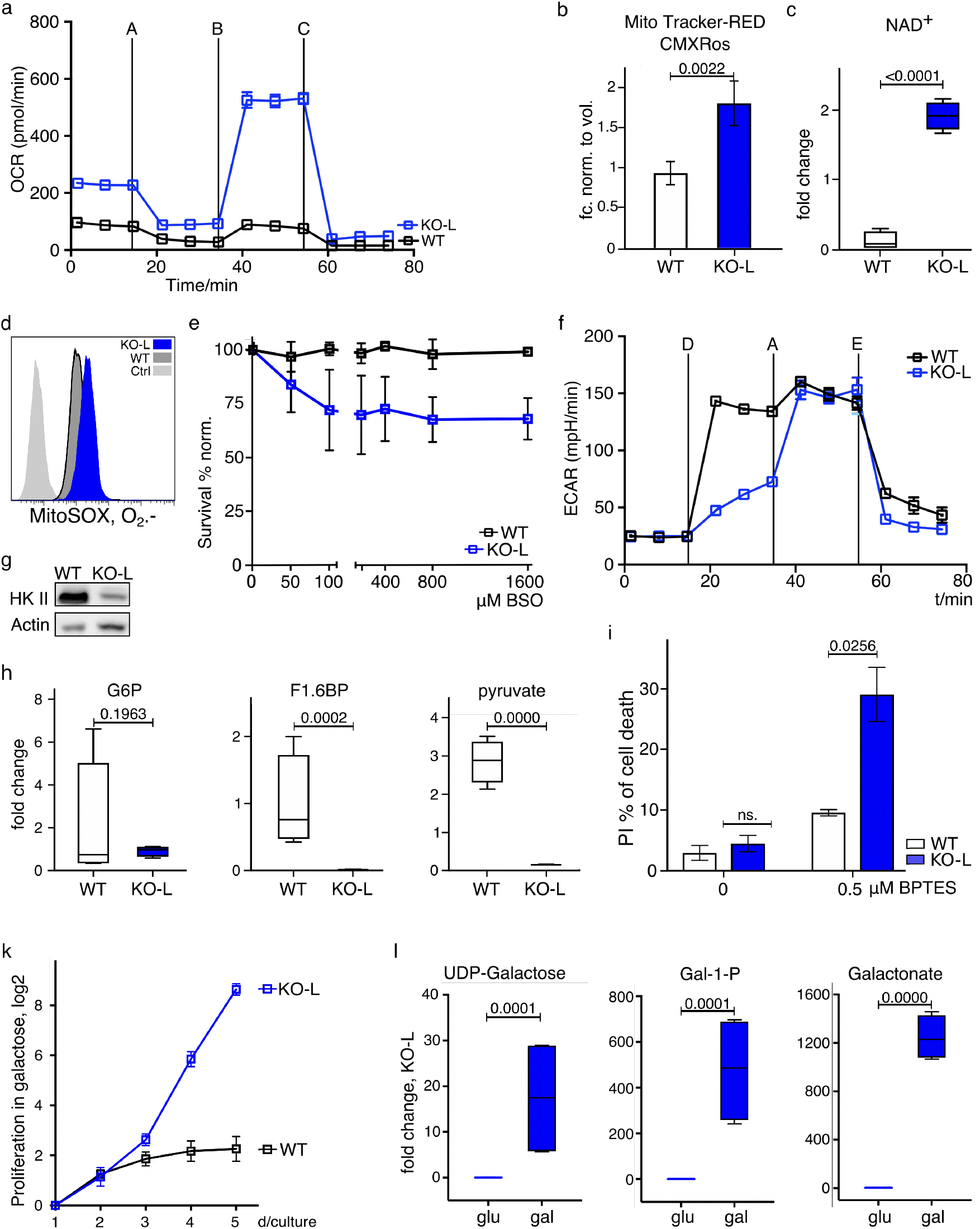
Metabolic switch of CD20KO Ramos B cells. **a** Oxygen consumption rate (OCR) of KO-L compared to WT Ramos cells. Metabolite flux analysis was performed in triplicates and is displayed as mean ± SD. One, out of 3 experiments, is shown. Used inhibitors: A= Oligomycin, B= FCCP, C= Rotenone + Antimycin. **b** Mitochondrial mass staining with Mito Tracker Red-CMXRos in KO-L compared to WT Ramos cells and normalized to volume of WT Ramos cells. n=4. **c** NAD^+^ levels of KO-L compared to WT Ramos cells determined with untargeted metabolomic profiling, n=3. **d** Mitochondrial superoxide levels stained with MitoSox Red. KO-L cells, WT Ramos cells and unstained control (Ctrl) are shown, n=4. **e** Survival rates (propidium iodide negative cells) of KO-L and WT Ramos cells after Buthionine sulfoximine (BSO) titration. The experiment was performed three times and mean MFI values were normalized to day 0. **f** Extracellular acidification rate (ECAR) of KO-L and WT Ramos cells as a measure of glycolysis is shown. The measurement was performed in technical triplicates and is displayed as mean ± SD. One, out of 3 independent experiments, is shown D= Glucose, A= Oligomycin, E= 2DG. **g** Representative western blot analysis of Hexokinase II (HK II) protein levels of KO-L and WT Ramos cells. Lysates were taken 20 days after induction of CD20 KO and data are representative for at least three independently generated CD20 KO Ramos B cell lines. **h** Levels of the glycolytic intermediates glucose 6-phosphate (G6P), fructose 6-phosphate (F1,6BP), and pyruvate as determined by untargeted metabolomic profiling are shown, n=3. **i** Cell death as determined by propidium iodide (PI) staining after treatment with BPTES, n=3. **k** Proliferation index determined with MFI of CellTrace staining per day (d) of KO-L cells cultivated in galactose compared to WT Ramos cells, n=3. **l** Levels of UDP-galactose, galactose-1-phosphate (G 1-P) and galactonate as determined by untargeted metabolomic, n=4.

Nicotine amid adenine dinucleotide (NAD+) is a crucial cofactor for enzymes in several metabolic pathways and high levels of NAD+ have been suggested to favor mitochondrial biogenesis and function^25^. We found mitochondrial mass (**Fig 5b**) and NAD+ levels (**Fig 5c**) to be significantly increased in KO-L Ramos cells in comparison to WT cells. Since reactive oxygen species (ROS) are a natural by-product of respiratory activity, we hypothesized that ROS production might be altered in KO-L Ramos cells. Indeed, we found O_2_^-^ levels to be increased (**Fig 5d**) in KO-L Ramos cells. Moreover, we found the glutathione-cysteine ligase inhibitor buthionine sulfoximine (BSO) to induce cell death of KO-L Ramos cells, and less so in WT cells (**Fig 5e**). Glutathione-Cysteine Ligase is an important component of the Glutathione-dependent ROS scavenging pathway and protects cells from ROS induced cellular damage. Our results thus suggest that KO-L Ramos cells are exposed to increased levels of ROS and that adaptations to oxidative stress are crucial for their survival.

We next compared the glycolytic activity of WT and KO-L Ramos cells by measuring the extracellular acidification rate (ECAR). As expected for a Burkitt lymphoma, WT Ramos cells showed robust increase of ECAR after receiving glucose (**Fig 5f**). Upon oligomycin treatment, which is an inhibitor of mitochondrial ATPase, ECAR was further increased to compensate for the loss of mitochondria-derived ATP. In contrast to WT cells, KO-L cells showed low lactate production upon receiving glucose, yet ECAR levels were similar to those seen in WT after oligomycin treatment (**Fig 5f**). This suggests that although KO-L cells are capable of converting glucose to lactate this metabolic pathway is not favored under normal conditions. Consistent with their reduced glycolytic rates, the KO-L cells showed reduced expression of hexokinase II (HKII), the enzyme mediating the first step of glycolysis (**Fig 5g**). Furthermore, we found the levels of glycolytic intermediates such as glucose 6-phosphate, fructose 6-phosphate, and pyruvate to be decreased in KO-L cells (**Fig 5h**). Taken together these data suggest that glucose is not the primary source of energy in KO-L Ramos cells.

To assess whether alternative carbon sources play a more prominent role in KO-L Ramos cells, we treated the cells with Bis-2-(5-phenylacetamido-1,3,4-thiadiazol-2-yl)-ethyl sulphide (BPTES), an inhibitor of glutaminolysis. We found that while BPTES treatment induced cell death of KO-L cells, WT Ramos cells were less affected (**Fig 5i**). Furthermore, unlike WT Ramos, KO-L cells can be cultured in a medium where glucose is replaced by galactose (**Fig 5k**). KO-L cells are able to process galactose intracellularly and use it as an alternative carbon source (**Fig 5l**). Unlike glucose, galactose needs to be catabolized in the mitochondria to provide cells with energy. Thus, our results demonstrate that KO-L Ramos cells can survive entirely without glycolysis-derived ATP.

In summary, our metabolic analysis demonstrates that the loss of CD20 results in profound metabolic alterations which in many instances mirror the phenotype of plasma cells/plasma blasts^26,27^.

## Discussion

Macfarlane Burnet’s clonal selection theory provided the foundation for the explanation of adaptive immunity. The proposed selection process requires a strict separation between resting and activated B lymphocytes. We show here that CD20 is a gatekeeper of the resting B cell state. On resting mature B lymphocytes, the IgM-BCR and the IgD-BCR reside on the cell surface inside separated nanodomains with distinct lipid and protein compositions. Together with CD19, CD20 is a component of the IgD-class nanodomains. Upon antigen-dependent B cell activation, this nanoscale receptor organization is altered and CD19 together with the IgM-BCR forms a signaling synapse. We find that the exposure to anti-CD20 antibodies *in vitro* and the loss of CD20 also results in IgM/CD19 synapse formation, association to Syk/PI-3 kinase signaling and transient B cell activation by itself.

CD20 deficiency in mice and humans has been associated with reduced B cell signaling and T cell-independent immune responses rather than increased BCR signaling^28,29^. These differences may be due to compensation mechanism operating during B cell selection^30,31^. With the CRISPR/Cas9 technique it is feasible to establish a CD20 deficiency at the mature B cell stage without interference of the B cell selection process and to combine several gene KO for a gene function analysis. In this way, we could show that the transient B cell activation is associated with plasma cell differentiation of all our Ramos cell lines. With an unbiased GSE transcriptome analysis we found plasma cell-specific genes upregulated after combined RTX treatment in the transcriptome of about 85% of the investigated patients^22^. Disruption of the nanoscale organization of CD20 followed by development of plasma cells could explain the results of a study on immune thrombocytopenia patients after RTX medication which suggests that B cell depletion generates a milieu that promotes the differentiation and settlement of long-lived PC in the spleen^32,33^.

It is known that plasma cells improve metabolic flexibility^34,35^ to secure function and prolonged survival. Our analysis of the metabolic program of KO-L Ramos cells allowed us to identify metabolic vulnerabilities in these cells. Our finding that CD20KO Ramos cells switch from glycolysis to oxidative phosphorylation and from glucose to glutamine usage is in line with previous studies of plasma cell metabolism^24,25^. Importantly, the insight obtained from this metabolic profiling enabled us to identify BPTES and BSO as inhibitors reducing the viability of KO-L Ramos cells and these compounds provide a rationale to target these metabolic pathways in plasma cell pathologies. To find plasma cell characteristics and adaptations on CD20 negative cancer B cells make a successful treatment more challenging but enables new possibilities for patients that have relapsed after RTX medication. They may benefit from a treatment tailored to make use of the specific biological properties of plasma cells.

## Material and Methods

### Primary human naïve B cells

Primary naive B cells were obtained from fresh buffy coats provided by the Institute for Transfusion Medicine and Gene Therapy, ITG Freiburg. Peripheral blood mononuclear cells (PBMCs) were separated by Ficoll gradient centrifugation of whole blood and negatively selected using EasySep Human Naïve B Cell Isolation kit (StemCell). Prior to the experiments, primary naïve B cells were controlled for purity and rested overnight in complete RPMI 1640 supplemented with 5% FCS, 10 units/ml penicillin/streptomycin (Gibco), 20 mM HEPES (Gibco), and 50 mM β-mercaptoethanol (Sigma) at 37°C with 5% CO_2_. Experiments with primary human B cells were repeated at least three times. This study was approved by the Institutional Review Board of the University Freiburg (Ethical votes No 507/16 and 336/16).

### Cell culture

The human Burkitt lymphoma B cell-line Ramos was obtained from ATCC, Ramos (RA 1) (ATCC^®^ CRL-1596™). Ramos cells were cultured in RPMI 1640 medium (Gibco), supplemented with 5% FCS (Biochrom), 10 units/mL penicillin/streptomycin (Gibco), 20 mM HEPES (Gibco), and 50 mM β-mercaptoethanol (Sigma) at 37°C with 5% CO_2_. For glucose versus galactose metabolism analysis, Ramos B cells were cultured in RPMI 1640 medium without glucose (Gibco), supplemented with 11 mM sterile filtered glucose or galactose, respectively, 5% dialyzed FCS (Thermo Fisher), 10 units / mL penicillin / streptomycin (Gibco), 20 mM HEPES (Gibco), and 50 mM β-mercaptoethanol (Sigma). Cells were cultured at 37°C with 5% CO_2_.

### CRISPR/Cas9 knock out

CRISPR/Cas9 KO plasmids used in this study are listed in **Supplementary Table 1**. CRISPR/Cas9 deletion was either carried out using the Neon Transfection System (Invitrogen) to either deliver the Human MS4A1 KO plasmids (Santa Cruz) into the cells or the ribonucleoprotein according to the genome editing method of Integrated DNA Technology (IDT)^36^.

For plasmid delivery, 1.1 × 10^6^ cells were resuspended with 110 μL transfection medium containing 20 mM HEPES (Gibco) and 1.25% DMSO (Sigma) in RPMI medium and then mixed together with 4 μg of KO plasmid. The cells were then subjected to Neon Transfection System. Electroporation was performed in 100 μL NEON tips at 1350 V, 30 ms and a single pulse and recovered at 37°C and 5% CO_2_ without antibiotics for one day and then cultured in complete RPMI-medium.

For the IDT CRISPR-Cas9 approach equimolar amounts of Alt-R CRISPR-Cas9 crRNA and tracrRNA were annealed in IDTE buffer, combined with Cas9 endonuclease before complex formation with RNP. One million cells were then subjected to Neon Transfection System. Electroporation was performed in 10 μL NEON tips at 1350 V, with a single 30 ms pulse. The transfected cells were cultured at 37°C and 5% CO_2_ in complete RPMI medium. Inactivation of the target gene was verified by flow cytometry and/or western blotting.

### Flow cytometry analysis

For surface staining, 1–20 × 10^5^ cells were stained with antibodies in PBS supplemented with 0.5% BSA and 0.05% NaN_3_ on ice for 20 min, washed twice and then measured with a FACS Gallios (Beckman Coulter) For intracellular staining, the cells were fixed with 4% PFA, washed twice in PBS and permeabilized with 0.5% BSA containing 0.5% saponine for 60 min at room temperature. Data were exported in FCS-3.0 format and analyzed with FlowJo software (TreeStar). A BIO-RAD S3e cell sorter was used to select CD20-negative cell populations.

Flow cytometry analysis were performed more than three times from each of the eighteen independently generated CD20KO Ramos B cell lines. Detailed information about primary antibodies are listed in **Supplementary Table 2**.

### Lymphocyte and B-cell subpopulations phenotyping

Phenotyping of T-, B- and NK cells within the lymphocyte population was performed by a whole blood staining lyse-no-wash protocol (Optilyse B, Beckman-Coulter) using six colour flow cytometry with fluorochrome-conjugated antibodies as listed in **Supplementary Table 7 top**. Analysis of B-cell subpopulations was performed with a bulk lysis specimen of EDTA anticoagulated whole blood treated with ammonium chloride lysis buffer to lyse erythrocytes. After washing, a lyse-wash protocol (Optilyse B, Beckman-Coulter) was performed. To determine B-cell subpopulations, nine colour flow cytometry with fluorochrome-conjugated antibodies as listed in **Supplementary Table 7 bottom** was performed. Gating strategy in **Extended Data Figure 4b**. Antibody labelled cells were analyzed by flow cytometry (Navios; Beckman Coulter). Flow cytometric data analysis was performed with the help of Kaluza Software 1.5a (Beckman Coulter).

### Cell survival

Cell survival was determined by analyzing forward/sideward scatter properties of cells and simultaneous staining with either Propidium iodide (PI), or 7-amino-Actinomycin D (7AAD eFluor670 dye, and LIVE/DEAD fixable violet stain (ThermoFisher), following the manufacturer’s instructions. Cell survival was continuously monitored in all experiments.

### Cell proliferation assay

The cell proliferation assay was performed using CellTrace Proliferation Kit (ThermoFisher) according to the manufacturer’s protocol. In brief, one million cells were washed with PBS and stained in a 1/1000 dilution of the stock solution for 20 min at 37°C protected from light. Cells were washed twice with complete supplemented RPMI. After 10 min the mean fluorescence intensity (MFI) was measured. Further readings were followed daily. MFI values were used to calculate the proliferation index^37^. Proliferation was measured at least three times.

### Western Blotting

Cells were collected and immediately lysed in 2x Laemmli or RIPA buffer. Equal amounts of cleared lysates were subjected to SDS-PAGE on 10%, 12% or gradient (10-15%) mini precast gels (7bioscience) and to subsequent immunoblotting on PVDF membrane (GE Healthcare Amersham) and blocked with 5% BSA in PBS and 0.1% Tween20. A horseradish (HRP) peroxidase-conjugated goat anti-rabbit or goat anti-mouse antibody was used as secondary antibody before detection with ECL chemiluminescent substrate (Bio-Rad). Western blot experiments were repeated more than three times. Lysates were taken from independently generated Ramos CD20KO B cell lines. Details of primary antibodies are shown in **Supplementary Table 3**.

### CRISPR riboswitch design and cKO Ramos cell generation

The CRISPR riboswitch was designed based on a published exon-skipping RNA device^13^. A minimized portion of the original, synthetic construct including part of the 5’ intron, the 3’ splice site, a tetracycline aptamer, the poised exon, the 5’ splice site, and part of the 3’ intron was targeted to intronic sequences of the human CD20 gene. The tetracycline aptamers C1 and C2 used here correspond to those reported in the M1 and M2 minigene constructs^13^. The intronic CRISPR guides were designed to avoid regulatory elements using the UCSC Genome Browser. The Alt-R^^®^^ Cas9 ribonucleoprotein complexes were assembled as mentioned above^36^. The riboswitch was amplified by PCR (CloneAmp, Takara; according to the manufacturer’s instructions) verified by agarose gel, and cleaned using a PCR purification kit. Details of primers used for the ex34 or ex45 homology directed repair (HDR) are listed in **Supplementary Table 1**. The HDR templates were mixed at fourfold molar excess with a single-strand DNA binding protein (ET SSB, NEB, M2401S) and heated at 95°C for 10 minutes to achieve singlestranded donor oligonucleotides (ssODNs) bound to ET SSB. About 2 pmoles of HDR template were added to the RNP-enhancer-cell mixture and subjected to Neon Transfection System. Electroporation was performed in 10 μL NEON tips at 1450 V with a single 20 ms pulse. For analysis four independent cKO Ramos B cells lines were created and Tet-induction experiments were performed at least four times.

### Surfaceome analysis using Cell Surface capture (CSC)

For surfaceome screening 50 million cells per replicate of Ramos WT, CD20KO-N and CD20KO-L were harvested and washed with ice-cold PBS. CSC was performed as previously described^23^. Deamidated peptides derived from N-glycosylated cell surface-residing receptors were analysed on a Q Exactive plus HF mass spectrometer (QE-HF MS) Thermo Scientific) coupled to an EASY-nLC 1200 instrument. The QE-HF MS was operated in positive ion mode and peptides were separated by reverse-phase chromatography on a 15 cm column in-house packed with ReproSil-Pur 120A C18-AQ 1.9 μm (Dr. Maisch GmbH), primed with 100% buffer A (99% H2O, 01% formic acid) and with a 70-minute gradient from 6 to 44% buffer B (99.9% acetonitrile, 0.1% formic acid) prior to injection. Mass spectrometry data were acquired in data-dependent acquisition (DDA) mode (TOP10). MS1 scans were acquired with 60,000 resolution, AGC target of 3 x 10^6^, IT of 15 ms and followed by high-energy collision dissociation (HCD) at 28%. MS2 scans were recorded with 15,000 resolution, AGC target of 1 x 10^5^ and IT of 110 ms. Raw files were analysed using the Trans-Proteomic Pipeline (v4.6.2) by searching with COMET (v27.0) against Uniprot KB (Swiss-Prot, Homo sapiens, retrieved April 2018). Carbamidomethylation was set as a fixed modification for cysteine, oxidation of methionine and deamidation of arginine were set as variable modifications. Peptide feature intensities of identified peptides with FDR ≤ 1%, presence of consensus NXS/T sequence and deamidation (+0.98 Da) at asparagines were extracted with Progenesis QI (v4.0, Nonlinear Dynamics) for label-free quantification. Statistical analysis was performed with MSstats (v 3.8.6) using iRT peptides to normalize across runs. Significance of differentially regulated proteins was determined by the threshold |fold-change| >1.5 and p-value < 0.05 of a two-sided t-test with the appropriate degrees of freedom. Benjamini-Hochberg method was used to account for multiple testing. Protein abundance changes and significance was visualized in volcano plots using R. Voronoi treemaps were generated using the FoamTree tool (https://carrotsearch.com/foamtree/). Hierarchical classification was created based on the functional annotation of human plasma membrane proteins by Almén et al.^38^. CSC experiments were performed in triplicates per condition. Acquired raw data will be deposited to PRIDE. MS experiment details and link are shown in **Supplementary Table 6**. The data (ID:PXD019874) will be public on PRIDE, as soon as the paper is accepted. For review use the link: http://massive.ucsd.edu/ProteoSAFe/QueryPXD?id=PXD019874 Username: MSV000085605 Password: KathrinKlaesener Once the data are public the link will be: ftp://massive.ucsd.edu/MSV000085605

### Proximity ligation assay (PLA)

For Fab-PLA, the PLA-probes were prepared as previously described^39^. In brief, F(ab)-fragments were prepared from the corresponding antibodies using the Pierce Fab Micro Preparation Kit (Thermo Fisher) according to the manufacturer’s protocol. In brief, after buffer exchange (Zeba™ spin desalting columns, Thermo Fisher) F(ab)-fragments were coupled with PLA probemaker Plus or Minus oligonucleotides according to the manufacturer’s protocol (Sigma-Aldrich) to generate Fab-PLA probes. For *in situ* PLA experiments, the cells were allowed to attach to polytetrafluoroethylene (PTFE)-coated slides (ThermoFisher) for 30 min at 37°C. Depending on the experiment, the cells were activated with 1mM freshly prepared pervanadate or treated with RTX and then fixed for 20 min with 4% paraformaldehyde in PBS. PLA was performed as described earlier^39^. PLA experiments were repeated at least three times, each in two technical replicates. Details about primary antibodies used for PLA-probes are provided in **Supplementary Table 4**.

### Imaging, Image analysis, and Data processing

All microscope images were acquired using Leica DMi8 microscope equipped with a 63 × oil immersion objective lens. For each sample, several images of at least 1000 cells were imaged from randomly chosen regions. All recorded images were analysed with CellProfiler 3.0.0. Raw data produced by CellProfiler were exported to Prism software (GraphPad, La Jolla, CA). The mean PLA signal count per cell was calculated from the corresponding images and presented as scatter dot plots with mean and standard deviation (SD).

### Statistical analysis

Nonparametric Mann-Whitney U-test was used for all experiments shown except results obtained from the metabolome analysis. Statistical analysis of our metabolomics data was performed by Metabolon Inc and Welch’s two samples t-test was used to determine statistical significance.

### ELISA of IgA-Secretion

Quantification of total IgA antibody secretion was measured by sandwich ELISA (Thermo scientific) with minor modifications. In brief, after centrifugation media aliquots of three serial dilutions (n=3) of culture supernatants of KO-L and WT Ramos cells were added to 96-well plates coated with 10 μg/mL of human anti lambda specific antibody and kept o/n at 4°C to calculate maximal effective concentration. RPMI and pure PBS served as a control. After extensive washes with wash buffer containing PBS, supplemented with 0.05% Tween20 and 0.01 NaN_3_, non-specific binding was blocked using PBS, supplemented with 1% BSA and 0.01% NaN_3_. Binding was revealed using biotinylated anti-IgA secondary antibody followed by Streptavadin-Horseradish Peroxidase, developed with 3,3’,5,5’-tetramethylbenzidine (TMB), and detected with Elisa microplate reader before normalization. Independently generated Ramos CD20KO-L cell lines were chosen for three repetitions. Details in **Supplementary Table 4**.

### Inhibitors

For inhibition Ramos cells were treated with Bis-2-(5-phenylacetamido-1,3,4-thiadiazol-2-yl)ethyl sulfide (BPTES, final concentration of 0.5μM) or L-Buthionine Sulfoximine (BSO, final concentration 0 – 1600 μM, all Selleck-Chemicals). Inhibitors were diluted in RPMI culture medium and treatment was performed as described in manufacturer’s protocol.

### Rituximab (RTX) treatment

RTX was kindly provided by F. Hoffmann-La Roche-AG. For Ramos cell treatment RTX was diluted to a final concentration of 10 μg/mL and performed at least three times.

### Transcriptome analysis

Massive Analyses of cDNA Ends (MACE-Seq) was performed by GenXPro GmbH in Frankfurt am Main using the MACE-Seq kit according to the manual of the manufacturers^40^. Briefly, cDNA was generated with barcoded poly-A primers during reverse transcription. The cDNA was fragmented and a second adapter was ligated. Competitive amplification was used to produce a library that was sequenced on an Illumina NextSeq500 machine. The reads were cleaned from adapter-residues and homopolymer stretches and annotated to the human genome (HG19) and read counts per gene was quantified. Differential gene expression was determined using the “DE-Seq2” package for p-value- and FDR calculation. (Technical replicates of Ramos WT n=3, biological replicates of independently generated CD20KO cell lines: KO-I n=4, and KO-L, n=5)

Differentially regulated genes between the different groups were identified using a linear model-based approach (limma R package^41^). Briefly genes with non-null read count in at least 2 samples were selected. After applying TMM normalization, a regular fitting method with unpaired design was used. Finally, genes with missing entrez IDs were filtered out for further analysis. Ajdusted p-value <0.05 was considered as significant.

### GEO data

Microarray gene expression raw-data from 13 CLL patients treated with rituximab^22^ were downloaded from GEO (GSE37168). CEL files from samples before treatment and after relapse were RMA normalized using the oligo R package. Enrichment of plasma cell differentiation up-regulated genes was performed using the fgsea R package^42^. Genes were ranked according to their Log FC and used as input in fgsea. Genes for the individual patients were ranked according to the Log2 FC comparing the expression at relapse against before treatment from GSE37168. Plasma up-regulated genes are highlighted by the ticks on x-axis in fgsea, where color code represents the enrichment score during the random walk over the ranked list of genes. Statistical significance was assessed using 1000 permutations where p-value <0.05 was considered as significant. Plasma cell specific gene set is listed in **Supplementary Table 8**.

### Detection of mitochondrial mass

To stain for mitochondrial mass, 10^5^ cells were stained in 100 μL 60nM Mito Tracker-RedCMXRos (Thermo Fisher) in Ramos medium for 30min at 37°C. Cells were washed twice before measurement. Cells were analysed using a flow cytometer Gallios (Beckman Coulter). Data were normalized according to cell volume. Three independent experiments have been performed.

### Detection of mitochondrial ROS

Detection of superoxide in mitochondria was performed with MitoSOX-Red mitochondrial superoxide indicator (molecular probes) according to the manufacturer’s protocol. In brief, 10^5^ Ramos cells were loaded with 1mL of reagent working solution (5μM MitoSOX in HBSS-buffer) and incubated in the dark for 10 min in at 37°C. The cells were washed three times with warm buffer and immediately subjected to imaging. Three replicate measurements were taken.

### Metabolomic analysis

For metabolomic profiling, cells were washed with PBS and snap frozen in liquid nitrogen. Samples were processed and analyzed by Metabolon Inc. The extracts were divided into five fractions: two for analysis by two separate reverse phase (RP)/UPLC-MS/MS methods with positive ion mode electrospray ionization (ESI), one for analysis by RP/UPLC-MS/MS with negative ion mode ESI, one for analysis by Hydrophylic interaction chromatography (HILIC)/UPLC-MS/MS with negative ion mode ESI, and one sample was reserved for backup. Raw data was extracted and compounds were peak identified by Metabolon Inc. Missing values were imputed with the observed minimum after normalization. Global metabolic profiles were determined using UPLC-MS/MS (Metabolon). Shown are cell count normalized results for NAD^+^, G6P, F1.6BP, pyruvate, UDP-Galactose, Gal-1-P, Galactonate. The raw data obtained from our metabolome studies will be deposited at MetaboLights upon manuscript acceptance.

### Metabolomic flux analysis

To measure glycolytic flux, cells were resuspended in 50μl Seahorse XF Base Medium (Agilent) supplemented with 2mM L-glutamine (Thermo Fisher) and incubated for 30min at 37°C in a CO_2_ -free incubator. Subsequently, 130 μL medium were added and cells were incubated for 1h. ECAR was measured using the Seahorse XFe96 metabolic flux analyzer (Agilent). Cells were sequentially treated with 10mM glucose (Sigma), 1μM oligomycin (Agilent), and 30mM 2-deoxy-D glucose (2DG) (Sigma). Oligomycin is an inhibitor of mitochondrial ATPse and 2DG inhibits the glycolytic enzyme hexokinase II. To assess mitochondrial function, cells were resuspended in 50μL Seahorse XF Base Medium (Agilent) supplemented with 2mM L-glutamine (Thermo Fisher), 1mM sodium pyruvate (Thermo Fisher), 10mM glucose (Sigma) and incubated for 30min at 37°C in a CO_2_ -free incubator. Subsequently, 130μL medium were added and cells were incubated for an additional 1h. Cells were sequentially treated with 1μM oligomycin (Agilent), 1μM FCCP (Agilent) and 1μM rotenone+antimycin (Agilent). OCR was measured using the Seahorse XFe96 metabolic flux analyzer (Agilent).

## Acknowledgments

We would like to thank Dr. Lise Leclercq for critical reading of the manuscript. We thank Qusai Al-Kayyal and Rita Rzepka for technical assistance. Many thanks to Marc Vogel for help and feedback concerning conditional riboswitch. Research in Wollscheid laboratory is funded by the ETH (grant ETH-30 17-1 to B.W.) and the Swiss National Science Foundation (grant 31003A_160259 for B.W.) This project was funded by the German research foundation (DFG) through TRR130 (project2 to M.R. project 25 to J.J. and project 12 to R.E.V.). Part of this work was financially supported by the Deutsche Forschungsgemeinschaft (DFG) SFB 850 and SFB1160 Z1 to MB. Additional support was received from the German Federal Ministry of Education and Research (BMBF) within the framework of the e:Med research and funding concept CoNfirm (FKZ 01ZX1708F to MB) and by MIRACUM within the Medical Informatics Funding Scheme (FKZ 01ZZ1801B to MB). Part of this work was supported by Roche, we thank Dr. Christian Klein, Roche Innovation Center Zurich for providing materials.

## Author contributions

K.K. and J.J. performed experiments, and analysed data. G.A. conducted mRNA transcriptome profiling. U.S. acquired patient-data. C.B. performed western blot experiments. M.C. designed CRISPR riboswitch and generated cKO cell-lines, J.B.A. and S.N.S. performed CSC with biostatistical analysis, B.W. provided expertise in proteotype and surfaceome analysis and acquired funding. B.S. sent material for gene editing of MS4A1 gene. R.E.V. provided expertise on primary B cells and buffy coats, M.B. enabled mRNA transcriptome analysis and funding, and M.R. secured funding, developed the concept and wrote the manuscript with the help of K.K., and J.J. All co-authors reviewed and edited the manuscript.

---

**Extended Figure 1:**
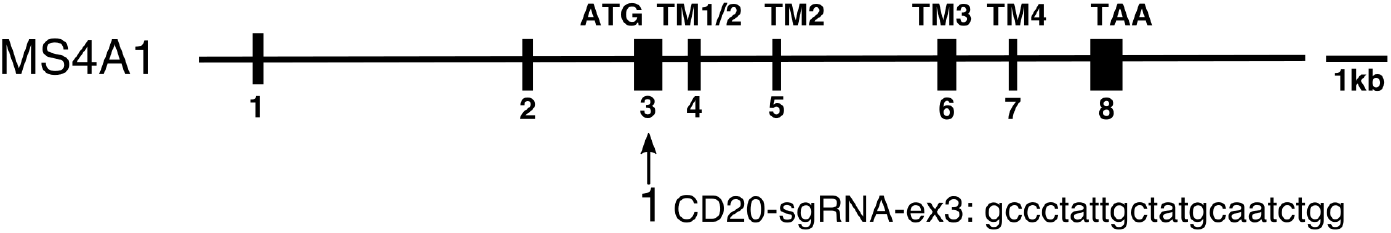
CRISPR/Cas9 CD20KO generation. Schematic of CRISPR-mediated gene targeting site in exon 3 of the scaled exon-intron organization of the human MS4A1 gene. Sequence of CRISPR-single guide RNA (sgRNA) as depicted.

**Extended Figure 2:**
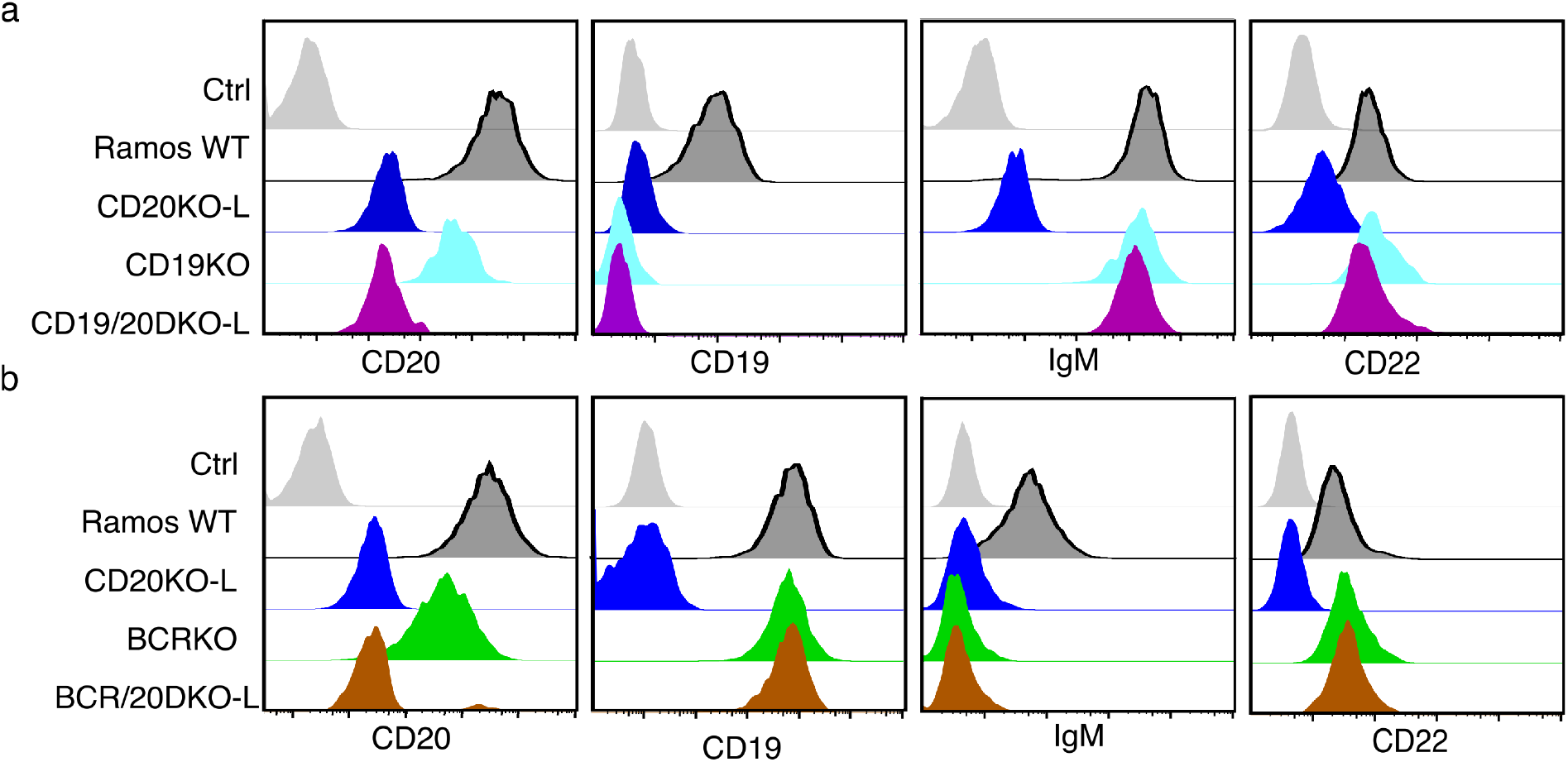
The IgM-BCR and CD19 are indispensable for CD20KO B cell activation. Flow cytometry analysis showing surface molecule expression on **a** CD19/20DKO-L, CD19KO, and CD20KO-L Ramos cells compared to WT and unstained control (grey) or **b** on BCR/20DKO-L, BCRKO, CD20KO-L Ramos cells compared to WT and unstained control (in grey).

**Extended Figure 3:**
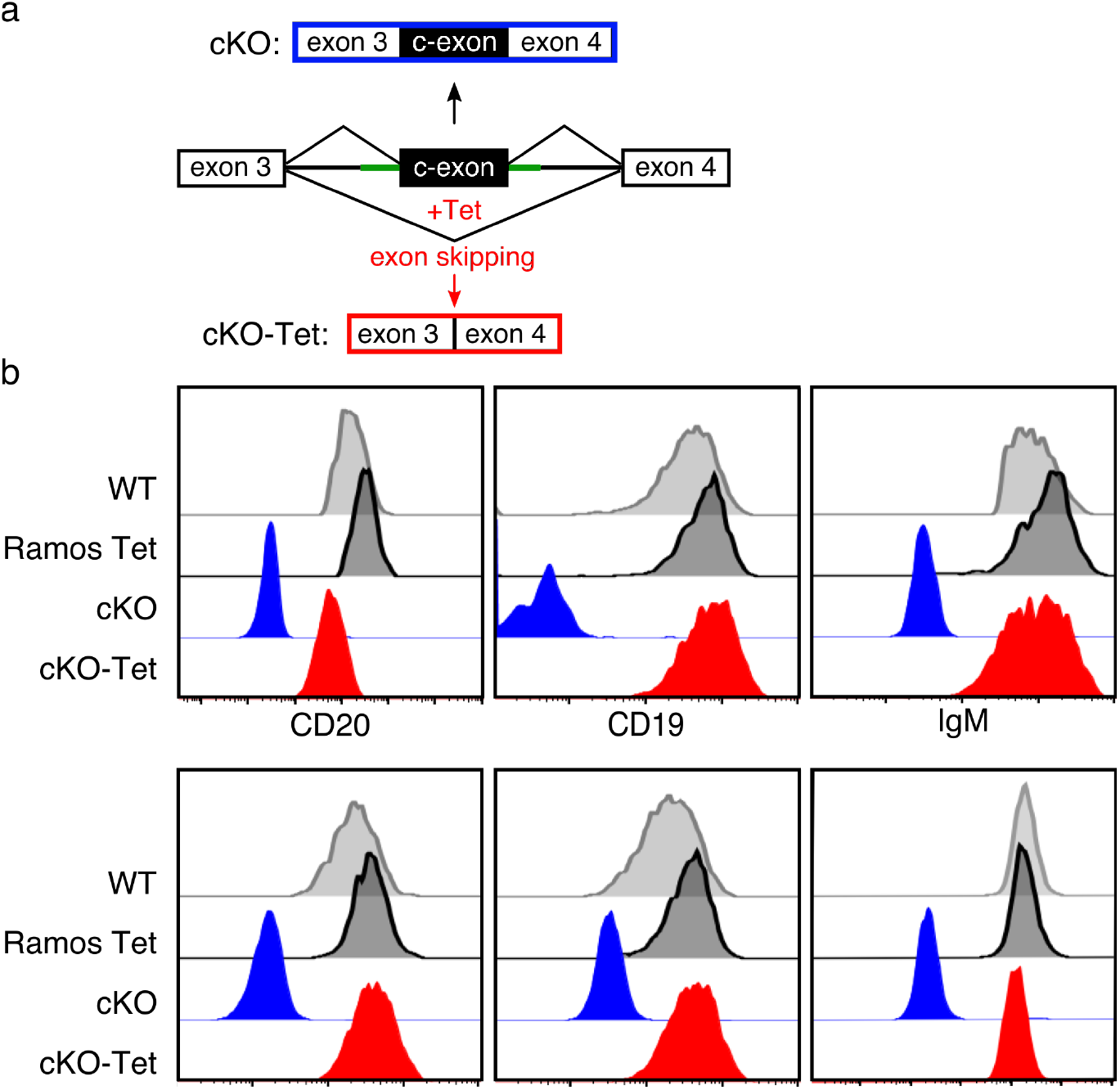
Proof of the Principle: conditional CD20KO (cKO) **a** Schematic of CRISPR-mediated insertion of an aptamer-controlled exon (c-exon) in intron 3-4 in the human MS4A1 gene generating a CD20 conditional KO (cKO-top). Tetracyclin (Tet) induces c-exon skipping and restores the ORF of the MS4A1 gene (cKO-Tet, bottom). **b** Induction of cKO Ramos B cells with 6μM Tet for 12h 30 days after transfection restored the WT phenotype. Examples of flow cytometry analysis of surface receptors CD20, CD19, IgM, CD22, CD81, and CD40 of cKO-Tet (red) compared to cKO Ramos cells (blue), WT and Tet-induced WT Ramos cells (Ramos-Tet) as control.

**Extended Figure 4.**
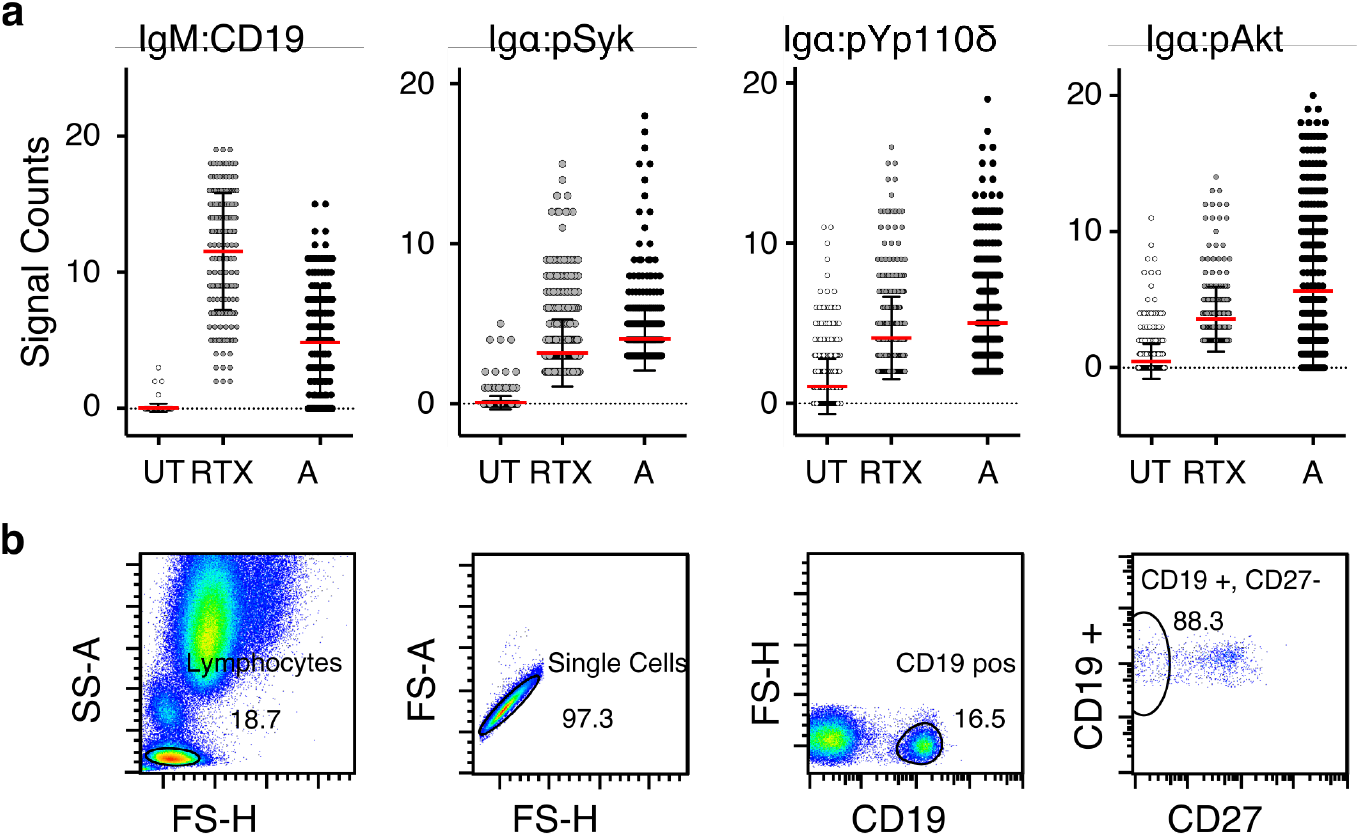
RTX treatment activates primary naïve B cells via the canonical BCR pathway. **a** PLA on resting (UT), 15 min RTX treated (RTX) or 5 min pervanadate activated (A) negatively selected naïve B cells from the peripheral blood of a healthy donor. Scatter dot plots showing the mean and SD of quantified results of the proximity of CD19 to IgM-BCR, Igα to phosphoSyk-Y525,526 (pSyk), Igα to phospho-p110δ-Y524 (pYp110δ), and of Igα to phosphoAkt-Ser473 (pAkt), n=4.

**Extended Figure 5.**
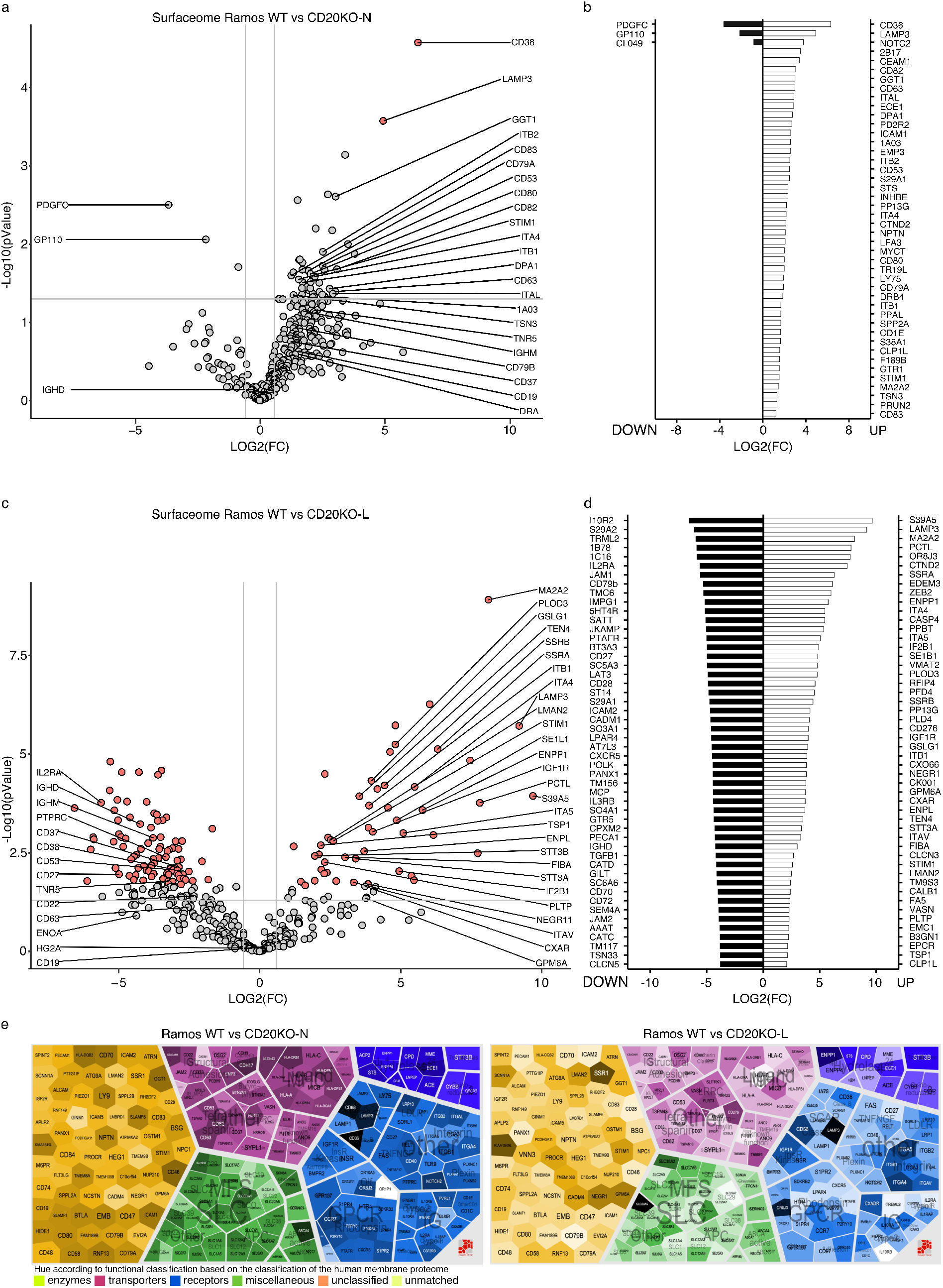
Surfaceome analysis of CD20KO-N and KO-L. **a** Volcano plot showing the Log2 FC of KO-N cell surface proteins compared to Ramos WT cells 10 days after gene targeting. **b** Top50 of significant (p-value < 0.05) of up (right) or downregulated (left) KO-N cell surface proteins compared to WT Ramos cells. **c** Volcano plot showing the Log2 FC of KO-L cell surface proteins compared to Ramos WT cells 1 month after gene targeting. **d** Top50 of significant (p-value < 0.05) of up (right) or downregulated (left) KO-L cell surface proteins compared to WT Ramos cells. **e** Voronoi tree map of KO-N surfaceome (left) or KO-L (right). All quantified proteins were mapped and hierarchically grouped according to their functional classification. For representation the Log2 FC values have been scaled from 0-1. The colors represent the Log2 FC between the compared conditions.

**Extended Figure 6.**
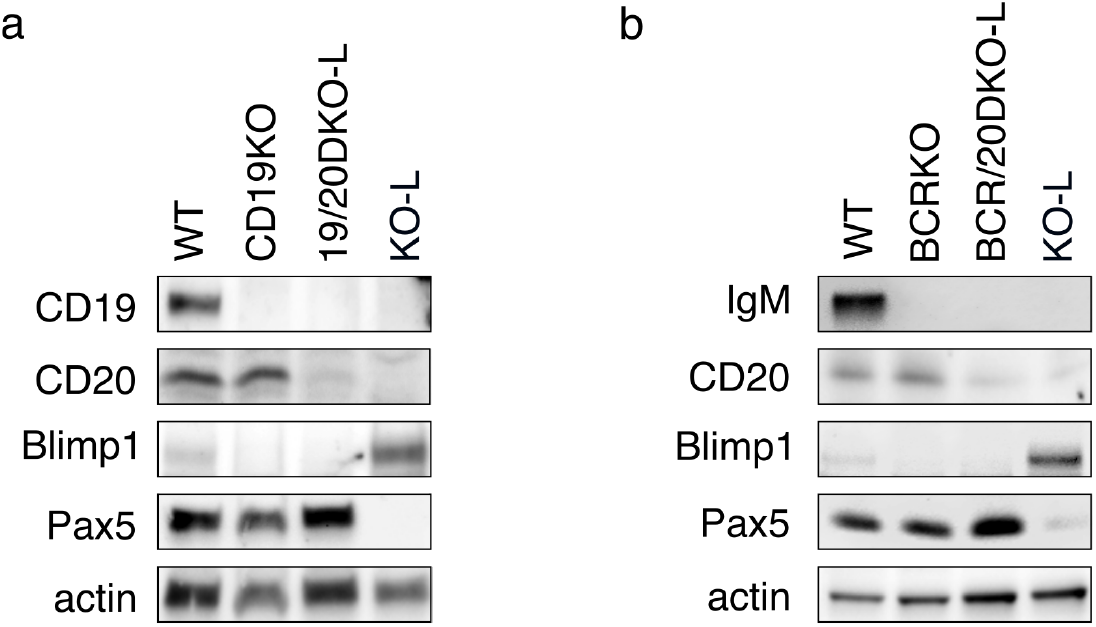
Expression of Pax5 and Blimp1 remain unchanged in CD19/20DKO and BCR/20DKO Ramos cells. **a** Representative examples of western blot analysis for B cell differentiation markers Pax5 and Blimp1 of CD19KO, CD19/20DKO-L, and KO-L Ramos B cells compared to WT. **b** Western blot analysis showing B cell differentiation markers Pax5 and Blimp1 of BCRKO, BCR/20DKO-L, and KO-L Ramos B cells compared to WT. All lysates were taken 20 days after induction of CD20 KO.

